# A biological growth curve is a sum of two distinct S-curves

**DOI:** 10.1101/2025.02.06.636984

**Authors:** I S Shruti

**Affiliations:** Independent Researcher, Alappuzha, Kerala, India

**Keywords:** growth curve, modeling, continued fraction

## Abstract

A growing population consists of successive generations of individuals interacting with each other and their environment. At every instant, these interactions can lead to either promotion or restriction of the population growth. Thus, within a growing population, differences arise over time due to both extrinsic and intrinsic factors such as resource utilization, competition, age distribution etc. These differences can lead to divergence within the population growth. The sum of these differences is manifested as the net population growth. In this work, we propose that these differences that arise over time within a growing population can be represented as two distinct S-curves. The sum of these two S-curves results in the growth curve of a population. These differences also arise in the growth curves of a population’s attributes such as height or weight. We demonstrate this by applying the *a*− *m* biological growth model on (i) the population growth curves of *Drosophila* and yeast and (ii) the growth curves of the mean height of a population of sunflower plants and the mean body weight of male white rats. Finally, we also discuss the results using a coordinate system with the two distinct S-curves as the axes.

## I Introduction

Based on his pioneering research on bacterial cultures, J. Monod described growth as a succession of phases characterized by variations in growth rate [1]. Although he has mentioned six phases, the order and presence of these phases can vary among different bacterial cultures. Since Monod’s classification of growth phases is based on the growth rate we generalize it to growth of all kinds. The phases are:

G1 lag phase (growth rate is null),

G2 acceleration phase (growth rate increases),

G3 exponential/linear phase (growth rate is constant),

G4 retardation phase (growth rate decreases),

G5 stationary phase (growth rate is null) and

G6 phase of decline (growth rate is negative).

In order to capture various phases, growth dynamics has been modeled as theory-based differential equations [2], [3], [4], empirical power laws [5] or polynomials [6], [7]. However, the phase of decline i.e. G6 has always been modeled separately [8].

In this work, we use a novel biological model that is based on the continued fraction of linear growth. This model, referred to as the *a* − *m* model has been shown to fit G2, G3 and G4 phases [9]. A linear combination of *a* − *m* models captures all the phases G1 to G6 of bacterial colonies at various antibiotic concentrations [13]. In the following section, we provide a brief overview of the *a*− *m* model. In Sec. III, we apply the linear combination model on *Drosophila*, yeast population data, height data of *Helianthus* and body weight data of male white rats.

## II. The *a* − *m* growth model

The *a m* model is obtained by introducing a parametric nonlinear term to the linear growth equation 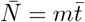.Thus, it is assumed that growth is fundamentally linear and the overall growth curve is a nonlinear deviation of linear dynamics at the G3 phase. The *a* − *m* model is given by

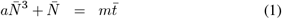

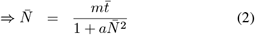

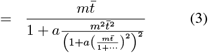

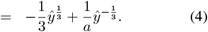

Eqn. (4) is obtained using Cardano’s method [12] for solving cubic equations (Eqn. (1)). Here, *ŷ* is given by

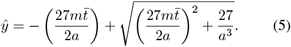

In the above equations, 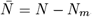,where *N* is the population size at time *t, N*_*m*_ is the population size at time *t*_*m*_ and 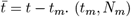 is the point of linear growth at G3 phase which is also the point of maximum growth rate *m*. When *N* = *N*_*m*_ and 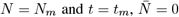 and 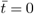,respectively. Therefore, the point (*t*_*m*_, *N*_*m*_) can be considered as the origin of the model. Here, nonlinearity is introduced through the parameter *a* as a continued fraction as in Eqn. (3).

The *a* − *m* model is actually a sum of two parts (Eqn. (4)). These two parts resemble power law relationships that are observed across biological systems [5]. They are:

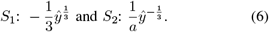

Therefore, the actual population size is given by

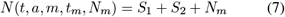

with the growth rate

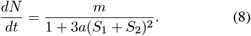

The *a* − *m* model is parameterized by two key factors: maximum growth rate *m* and a positive growth restricting term *a*. According to this model, the maximum growth rate is observed during the linear phase G3 when the non-linear restrictions of growth represented by *a* are minimal. This model was tested on population growth data of yeast cells and *Drosophila* as well as mean height and weight growth data of a population of sunflower plants and male white rats, respectively [9]. Taking the point of maximum growth rate as the origin, the *a* − *m* model was fit to the data such that it effectively captured the acceleration (G2), linear (G3) and deceleration (G4) phases of growth that appear in succession. The fit revealed that the G2 and G4 phases are distributed symmetrically around the G3 phase with respect to time. The *a*− *m* model was also fit on a historical growth (height) data of a human male provided by R. Scammon [10], [11]where it showed the same symmetry in the G2 and G4 phases as mentioned before. Moreover, it was also observed that even for periods as long as 6 years this symmetry was maintained (Fig. 7 of [9]).

In the following section, a weighted sum of *a*− *m* models is fit to the growth data of populations.

## III. Superposition of *a* − *m* models

As discussed in Sec. (I), the *a* − *m* model with the point of maximum growth (*t*_*m*_, *N*_*m*_) as the origin effectively fits the G2, G3 and G4 phases of growth.

In this section, we consider a weighted sum of multiple *a* − *m* models of different origins as a growth model. Such a model has been shown to fit all the phases of growth (G1 to G6) [13]. For a multi-origin model, the origins are chosen in the following manner [13]:

1. The absolute slope value between the *i*^*th*^ and (*i* + 1)^*th*^ data points is obtained as

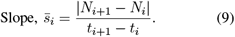
2. The first origin say, (*t*_*m*1_, *N*_*m*1_) is chosen as the midpoint of data points with the highest 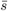.
3. The next origin (*t*_*m*2_, *N*_*m*2_) is the midpoint of data points with the second highest 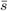.Subsequent origins are chosen in a similar manner.

This way multiple origins within the data points are selected. Thus, the final model is a weighted sum of *a* − *m* models (Eqn. 7) of different origins. Here, we use a model with three origins given by

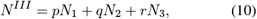

where *N*_1_ = *N* (*t, a, m*_1_, *t*_*m*1_, *N*_*m*1_), *N*_2_ = *N* (*t, a, m*_2_, *t*_*m*2_, *N*_*m*2_) and *N*_3_ = *N* (*t, a, m*_3_, *t*_*m*3_, *N*_*m*3_), where *N* is given by Eqn. (7). So, this model has 7 parameters *p, q, r, m*_1_, *m*_2_, *m*_3_ and *a*. Although this model has several parameters there is only one nonlinear parameter *a*. This is evident from the following form using Eqn. (2),

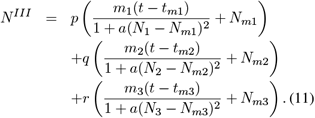

The above equation can be written in terms of *S*_1_ and *S*_2_ as

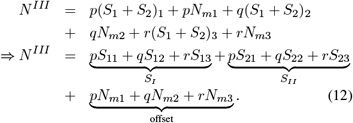

Therefore we have,

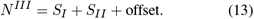

Thus, the final model is expressed as a sum of two S-curves with an offset. This model is fit to the growth data shown in Figs. (1) to (4) using the lmfit package [14] with initial guesses: *p, q, r* = 0; m_1_, *m*_2_, *m*_3_ = 1 and *a* = 1*e*− 3 with *a >* 1*e* − 6 for the standardized data.

**Fig. 1:**
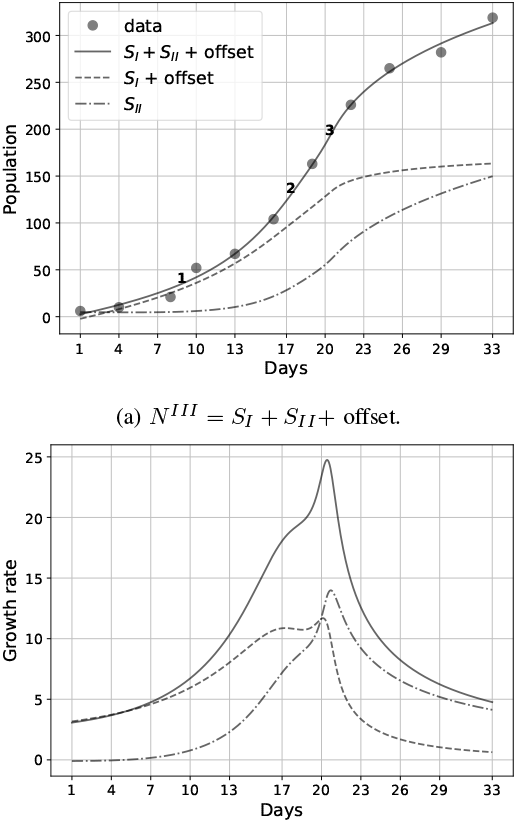
Successions in *Drosophila* growth data provided by Pearl [15]. *a* = 1.83*e* −5, *p* = −0.38, *q* = 0.66, *r* = 0.11, *m*_1_ = 8, *m*_2_ = 25.82 and *m*_3_ = 122.58. Origins ‘1’, ‘2’ and ‘3’ are also shown in (a).

The growth rate of *N*^*III*^ is given by

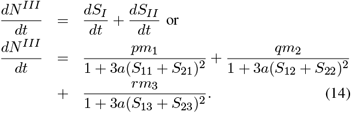

Therefore, *pm*_1_, *qm*_2_ and *rm*_3_ represent the maximum growth rate parameters of *a* − *m* models of different origins. The implementation of the fitting procedure is presented in Table (I). The growth rates can be obtained directly as follows:

**TABLE I:**
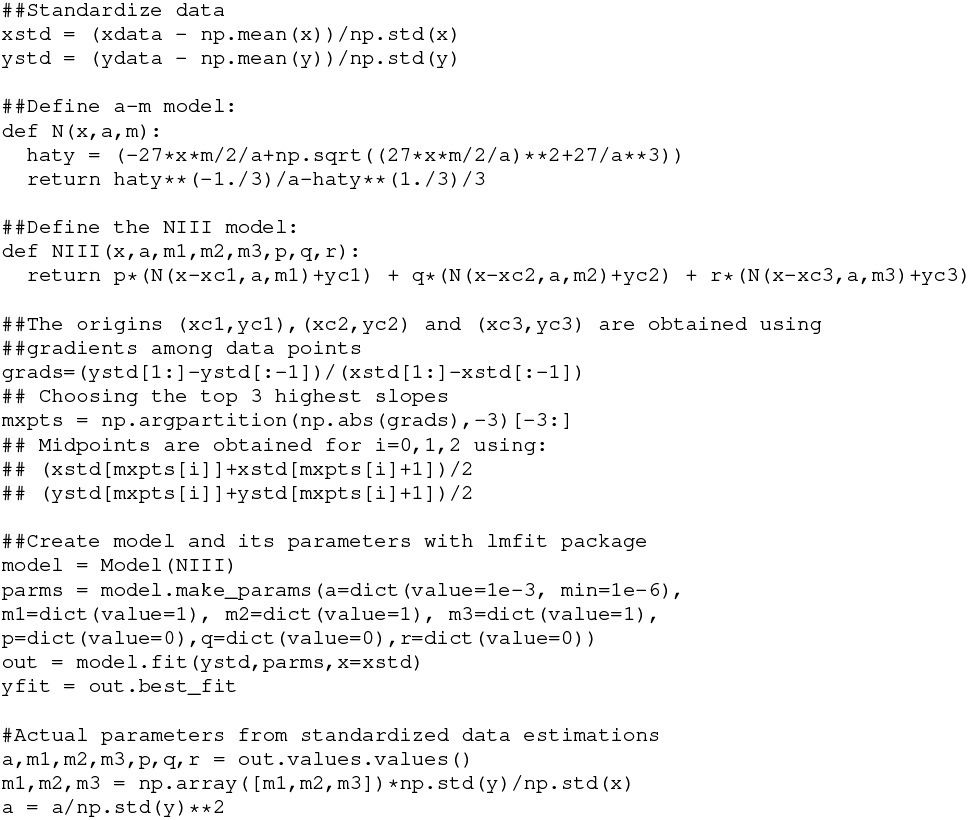
Python implementation of the fitting procedure.

~~~
def Ngrad(x,a,m):
    return m/(1+3*a*N(x,a,m)**2)
IIIgrad = p*Ngrad(x-xc1,a,m1)
+ q*Ngrad(x-xc2,a,m2)
+ r*Ngrad(x-xc3,a,m3)
~~~

We will now obtain values of parameters for various growth data below. For a linear model the maximum growth rate would be a weighted sum of the maximum growth rate parameters i.e. just the numerators of Eqn. (14). Therefore the linear growth rate is given as *m*_linear_ = *pm*_1_ + *qm*_2_ + *rm*_3_. However, the actual growth rate *m* would be the maximum value of Eqn. (14). Thus, the discrepancies between *m*_linear_ and *m* indicates the effect of nonlinearity through *a* on the maximum growth rate. The single-origin *a*− *m* model and the multi-origin models are compared in Table (II).

**TABLE II:** Summary of models based on the continued fraction of straight lines.

| One-origin [9] | Multi-origin [13] |
| --- | --- |
| 2- parametric | 5- (or more) parametric |
| $m$ is independent of $a$ | $m$ depends on $a$ |
| $m$ is directly estimated from the fitted curve | $m$ is the peak value of the growth rate curve |
| fits G2,G3 and G4 phases | fits all the phases G1 to G6 |
| Captures either growth or decline | Captures growth and decline simultaneously |
| Helps to establish symmetric nature of G2 and G4 | Provides a measure for nonlinearity |

### A. Drosophila Population

We now consider the *Drosophila* population data provided by Pearl (Table 11 of [15]). In an earlier study [9], an *a* − *m* model with a single origin (*t*_*m*_ = 19.44, *N*_*m*_ = 166.56) was fit on this growth data after excluding the first two data points. The maximum growth rate was found to be *m* = 23.92 for the selected data points.

In this work, we fit the *N*^*III*^ model on the growth data without omitting any of the data points as shown in Fig. (1a). The maximum growth rate parameters of *a* − *m* models with origins ‘1’, ‘2’ and ‘3’ (i.e., *N*_1_, *N*_2_ and *N*_3_ of Eqn. (10)) are obtained as *pm*_1_ = − 3.04, *qm*_2_ = 17.04 and *rm*_3_ = 13.48, respectively.

Since *pm*_1_ is negative, the corresponding *a*− *m* model *N*_1_ represents a decline in the number of individuals while *N*_2_ and *N*_3_ reflect the growth in number with time. The net maximum growth rate obtained from Fig. (1b) is *m* = 24.75. This maximum growth is less than the linear sum of *N*_1_, *N*_2_ and *N*_3_ growth rates which is: *m*_linear_ = *pm*_1_ + *qm*_2_ + *rm*_3_ = 27.59.

#### a) Role of a

As shown in Fig. (1), since both *q, r >* 0, the growth with rate *m*_linear_ = 27.59 is restricted positively to *m* = 24.75 through *a*. This disparity between *m* and *m*_linear_ indicates that *a* has a restricting effect on the linear growth which leads to the actual value of *m* = 24.75. This restriction occurs as a continued fraction (Eqn. (3)) which includes restrictions at all time scales.

In Fig. (1a), initially *S*_*II*_ is almost null while *S*_*I*_ grows. Therefore, *S*_*I*_ may represent the growth of initial generations of individuals involved in establishing a constantly growing population in a pristine environment. *S*_*II*_, on the other hand, may represent the growth of subsequent generations that become part of a well-established population yet an increasingly constrained environment. The offset given by Eqn. (12) is added to *S*_*I*_ for a better graphical representation. Here, the offset is added in such a way that the initial values of *S*_*I*_ lie along the *N*^*III*^ curve and the initial values of *S*_*II*_ are close to 0. In Fig. (1a), the growth of *S*_*II*_ accelerates when the growth of *S*_*I*_ starts to decline. This may be due to the disparities that arise within the population as younger individuals arrive and older ones die out. Thus the growth of a population is a combination of successive S-curves.

Fig. (1b) is given by Eqn. 14 which has no influence due to offset. From the figure, it is seen that the first generation *S*_*I*_ peaks before *N*^*III*^ does while *S*_*II*_ peaks after *N*^*III*^. This clearly indicates that growth occurs in successions. In the following section, we will extend this observation to the population growth of yeast cells.

### B. Yeast Population

Using the yeast population data provided by Pearl (Table 9 of [15]), we use the *N*^*III*^ model to obtain a good fit and the resulting plots are shown in Fig. (2). The maximum growth rate parameters of *N*_1_, *N*_2_ and *N*_3_ are given by *pm*_1_ = 170.33, *qm*_2_ = 192.16 and *rm*_3_ = − 252.48, respectively. Here, the negative value of *rm*_3_ corresponding to the *N*_3_ model indicates a decline in the number of yeast cells while *N*_1_ and *N*_2_ represent increase i.e. growth. The net maximum growth rate *m* is obtained from Fig. 2b as *m* = 94.5 which is again less than the linear sum *m*_linear_ = *pm*_1_ +*qm*_2_ +*rm*_3_ = 109.2. As mentioned in the previous section, the disparity between *m* and *m*_linear_ is decided by the restricting effect that *a* has on linear growth.

**Fig. 2:**
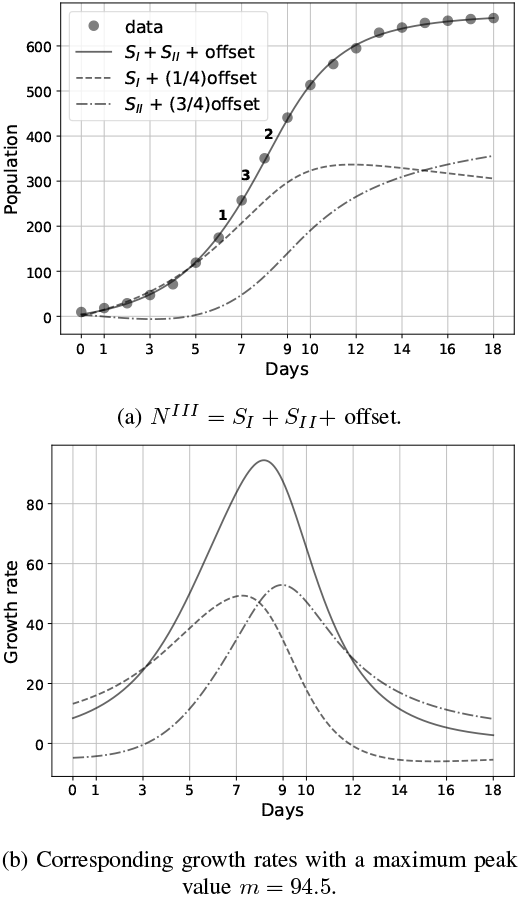
Successions in yeast growth data provided by Pearl [15]. *a* = 8.36*e* − 3, *p* = 138.48, *q* = 111.72, *r* = −277.45, *m*_1_ = 1.23, *m*_2_ = 1.72, *m*_3_ = 0.91.

Similar to the *Drosophila* plot, here too the *S*_*I*_ and *S*_*II*_ curves show successive growth. The *S*_*I*_ curve increases initially while *S*_*II*_ remains stagnant. However, when *S*_*I*_ starts declining, *S*_*II*_ continues to show accelerated growth. From Fig. (2b), it is observed that *S*_*I*_ peaks before *N*^*III*^ and *S*_*II*_ peaks after *N*^*III*^, thus indicating the shift in generations leading to the combined growth of the population. In the next two sections, we apply the *N*^*III*^ model on mean individual attributes such as height and weight.

### C. Height of Helianthus (sunflower) plants

The *Helianthus* growth data provided by Reed et al [16] consists of mean individual heights of fifty eight sunflower plants measured for eighty four days. Hence, here *N* represents the centimeter growth in height of *Helianthus* plants. Fig (3a) shows the fitting of *N*^*III*^ model on this growth data and the corresponding *S*_*I*_ and *S*_*II*_ curves. The maximum growth parameters of *N*_1_, *N*_2_ and *N*_3_ are *pm*_1_ = 28.57, *qm*_2_ = 15.33 and *rm*_3_ = − 36.3, respectively. *rm*_3_ being negative, we can conclude that *N*_3_ represents decline while *N*_1_ and *N*_2_ represent growth in height.

**Fig. 3:**
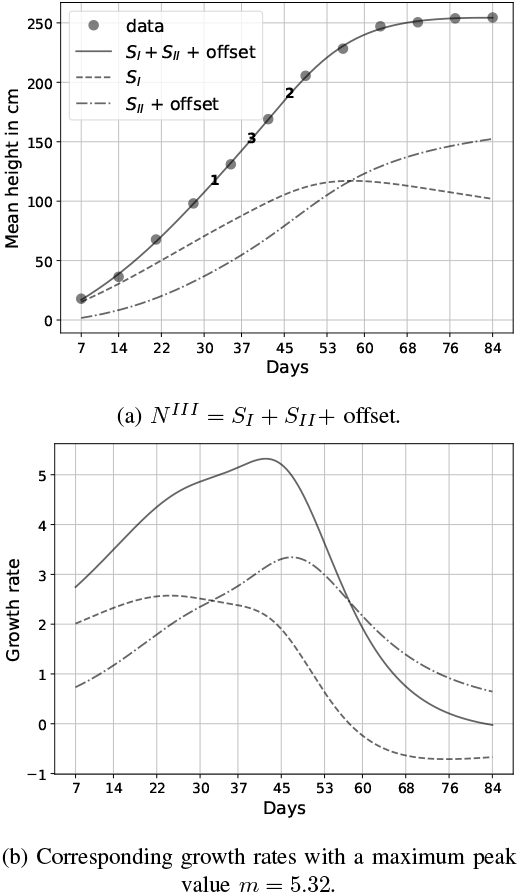
Successive growth in height of *Helianthus* plants based on the data provided by Reed et al [16]. *a* = 7.22*e* − 6, *p* = 4.77, *q* = 1.59, *r* = −6.6, *m*_1_ = 5.99, *m*_2_ = 9.64, *m*_3_ = 5.5

From Fig (3b), the actual maximum growth rate is obtained i.e. *m* = 5.32. This is considerably lower than the linear maximum growth rate given by *mlinear* = *pm*_1_ + *qm*_2_ + *rm*_3_ = 7.54. This indicates the highly nonlinear nature of this data as the restriction of linear growth through *a* is quite influential here. This is also reflected in the values of relative difference between *m*_*linear*_ and *m* given in Table (III). The *Helianthus* data shows the highest relative difference of 41.72% among all the data.

**TABLE III:** The nonlinearity through *a* is evaluated using the relative difference |*m*_linear_ − *m*|*/m*.

| Growth data | $\frac{\vert m_{linear} - m\vert }{m} \times 100$ |
| --- | --- |
| *Drosophila* | 11.5% |
| Yeast | 15.55% |
| *Helianthus* | 41.72% |
| Male white rats | 3.24% |

The *S*_*I*_ and *S*_*II*_ curves given in Fig. (3a) may represent two distinct spurts of growth in height of the plants. Also, *S*_*I*_*I* peaks much later than *S*_*I*_. This may be physically interpreted as some of the plants showing stagnation in growth, thus reducing the mean height of the data.

### D. Body weight of rats

The data considered here was provided by Donaldson [17]. Here *N* represents growth in body weight of male white rats measured in grams. As shown in Fig. (4a), the *N*^*III*^ model is fitted to the entire data and the maximum growth rate parameters are obtained as *pm*_1_ = − 5.36, *qm*_2_ = − 0.24 and *rm*_3_ = 8.01. Here, both *pm*_1_ and *pm*_2_ corresponding to *N*_1_ and *N*_2_ are negative. Hence, *N*_1_ and *N*_2_ represent decline in body weight which means that *a* also plays a role in restricting decline. The value of maximum growth rate *m* is obtained from Fig. 4b as 2.47. This is more than the linear growth rate given by *pm*_1_ + *qm*_2_ + *rm*_3_ = 2.39. This may be because the number of declining quantities that are being restricted is more than the growing quantity.

**Fig. 4:**
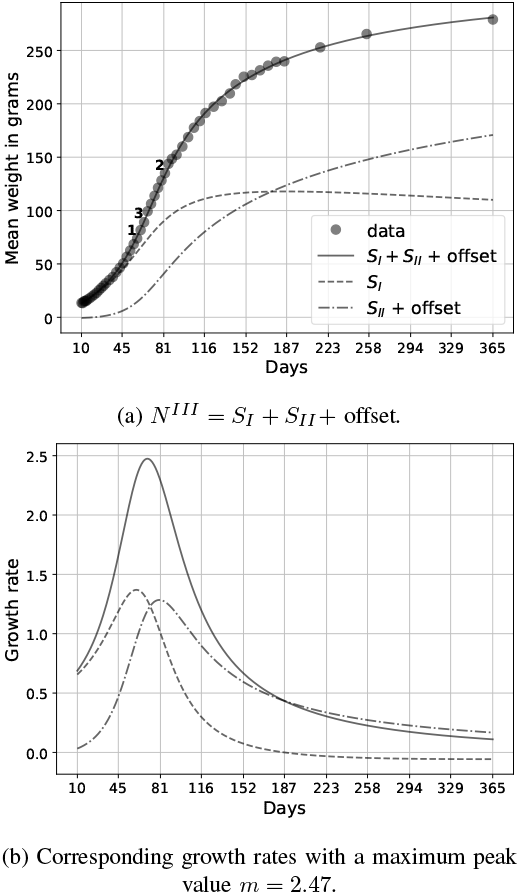
Successive growth in weight of male white rats based on the data provided by Donaldson [17]. *a* = 9.08*e* − 4, *p* = −17.74, *q* = −8.35, *r* = 21.65, *m*_1_ = 0.302, *m*_2_ = 0.029, *m*_3_ = 0.37

In Fig. (4a), it is seen that *S*_*I*_ first increases and then declines while *S*_*II*_ is almost null at first and then starts increasing gradually. The change in both curves is so gradual that it almost appears linear. This is also reflected in the relative difference between *m*_*linear*_ and *m* given in Table (III) as 3.2%. This is the least among all the data indicating that the nonlinear influence through *a* is minimal.

## IV. S_I_S_II_ PLANE

In this section, we present a coordinate system with the offsetted *S*_*I*_ and *S*_*II*_ as its axes. This is conceptually same as the *i, ii*− plane given in [18]. Fig. (5) summarizes the growth curves presented in the previous section. The offsets included in Fig. (5) are added in the same way as given in Figs. (1) to (4). In all the curves of Fig. (5), initially only *S*_*I*_ increases. Beyond a point, *S*_*I*_ either stagnates or starts to decline and *S*_*II*_ starts to grow. From Table III, it is seen that the growth in mean height of *Helianthus* plants is highly nonlinear. This is also seen in Fig. (5). In the *Helianthus* growth curve, *S*_*I*_ shows a sudden decline while *S*_*II*_ starts to grow. The yeast data shows a similar trend as the *Helianthus* data but shows a more gradual decline of *S*_*I*_. The rat data shows a gradual shift from *S*_*I*_ growth to *S*_*II*_ growth. The *Drosophila* growth data also shows a gradual shift yet not as smooth as the rat data.

**Fig. 5:**
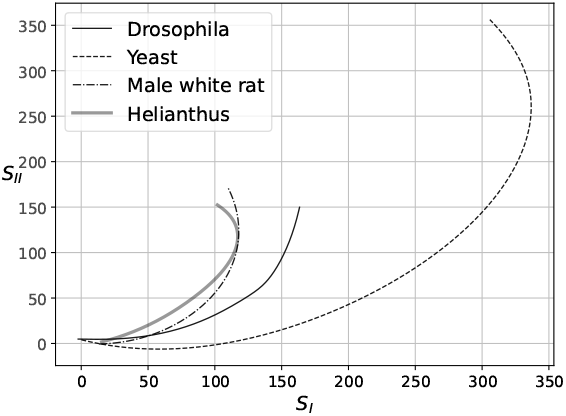
*S*_*I*_*S*_*II*_ plane including offsets from Figs. (1) to (4).

The *S*_*I*_*S*_*II*_ coordinate system can be considered as a bounded *xy*− plane [18]. This is because when *a* = 0 both *S*_*I*_ and *S*_*II*_ become unbounded and we get the original linear equation 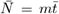.This is same as the linear equation *y* = *mx* on an *xy*− plane with *m* as the slope. Conventionally, we use *y* = *mx* and span the *xy*− plane with a single parameter *m*. Having an additional parameter *a* causes the dependent variable *y* to increase at a slower rate than the independent *x* values by following the path of an S-curve. Thus, *a* can also be considered as a bounding parameter that transforms unbounded straight lines in to S-curves.

## V. Conclusions

In this work, we have fitted the growth data of populations and individuals with a weighted sum of three *a*−*m* models of different origins and with the respective maximum growth or decline rate parameters. Thus, the weighted sum model includes growth and decline and shows a good fit for all the data. This means that the net growth of populations and individuals is a combination of growth and decline. The net growth curve is also shown to be the sum of two offset S-curves. It is also observed that the growth rate curves derived from these two S-curves peak in successions, and the curve of their sum i.e. the net growth rate peaks in between their peaks. The sum of the maximum growth rate parameters of the *a*− *m* model is considered as linear growth and is compared to the actual maximum growth rate obtained as the peak value of the net growth rate curve. From this, the relative difference between linear growth rate and the actual growth rate is calculated, which is helpful in analyzing the nonlinear dynamics of growth observed in these data. The growth data can be further visualized using a coordinate system with the offset S-curves as the axes. Since the growth data used here represent populations and individuals of different organisms, it can be concluded that biological growth is a combination of successive S-curves over time.

